# Phosphoproteomics in *Daphnia magna* as a tool to decipher molecular mechanisms in ecotoxicological studies

**DOI:** 10.64898/2026.05.01.721871

**Authors:** Magdalena V. Wilde, Jan B. Stöckl, Miwako Kösters, Marco M. Rupprecht, Julian Brehm, Michael Schwarzer, Kathrin A. Otte, Christian Laforsch, Thomas Fröhlich

## Abstract

Pollution of aquatic environments poses an increasingly severe threat to ecosystems worldwide, and understanding its molecular consequences for aquatic organisms requires extensive research and the development of advanced analytical tools. Phosphoproteomics can be particularly valuable for this purpose, as shifts in phosphorylation states can serve as early molecular indicators of toxic exposure. The cladoceran *Daphnia* is a keystone species in aquatic ecosystems, linking lower and higher trophic levels, and is therefore widely used as a model organism in ecotoxicology to study biological consequences of pollution. Here, we present a simple and effective strategy to analyse the phosphoproteome of *Daphnia magna*, a commonly used *Daphnia* species in ecotoxicology. Following TiO_2_-based phosphopeptide enrichment and LC-MS/MS analysis, we identified a comprehensive dataset of 3,532 phosphorylation sites across 1,329 phosphoproteins. These proteins were especially involved in signaling pathways and cellular structure and the vast majority have not yet been demonstrated in other *Daphnia* species. In conclusion, our results demonstrate that a straightforward phosphoproteomic LC-MS/MS workflow in *D. magna* can serve as a powerful tool for investigating adverse molecular effects caused by anthropogenic pollution, such as microplastics or pharmaceuticals.

**Statement of significance:** The dataset presented here demonstrates the feasibility of a simple yet effective strategy to perform phosphoprotemics in *Daphnia magna*, and it will be particularly valuable for future ecotoxicoproteomics research using this model organism.

---

Environmental pollution is one of the biggest challenges of our time. Therefore, a comprehensive understanding of the molecular responses underlying various pollutants (e.g. microplastics, pharmaceuticals, pesticides) is of utmost importance. One of the most important biochemical processes for modulation of protein activity in eukaryotic organisms are posttranslational modifications (PTMs) [1, 2]. Protein phosphorylations are reversible and very prominent PTMs and play a key role in intracellular signal transduction and affect nearly all basic cellular biochemical events, such as cell cycle, differentiation, and proliferation [3–5]. Changes in protein phosphorylation capture early pathway-level stress signaling and can therefore serve as early indicators of environmental change [6, 7]. Phosphoproteomics serves to identify and quantify protein phosphorylations throughout the proteome and provides a better understanding of complex phosphorylation-based signaling networks [4, 8, 9]. Understanding these phosphorylation-coordinated networks necessitates knowledge of specific amino acid modifications with both spatial and temporal resolution, which remains analytically complex and challenging [10]. For instance, phosphopeptides represent only a small fraction of all peptides present in a cell lysate due to the low stoichiometry of site-specific phosphorylations, and therefore need enrichment before LC-MS/MS measurements, requiring more sample material than standard analyses [4, 9]. Although several phosphoproteomic studies have been conducted in the field of ecotoxicology, improving the understanding of the effects of pollutants on organisms [11–14], to our knowledge, there has only been one study addressing the phosphoproteome of the cladoceran *Daphnia*. In 2014, Kwon et al. [15] performed the first global screening of phosphoproteins and their phosphorylation sites in *Daphnia pulex*. As *Daphnia* are key species of lake and pond ecosystems and are often used in ecotoxicological studies [16, 17], deeper insights into their molecular response to contaminants can help to understand the consequences of environmental pollution, especially since freshwater ecosystems suffer from various manmade threats (e.g., plastic pollution, agricultural activities, or sewage) [18]. Here, we present a comprehensive dataset of phosphorylated *D. magna* proteins that can serve as a reference for future proteomics studies on this organism, whether in an ecotoxicological context or to identify fundamental molecular processes.

Implementing an easy-to-perform phosphoproteomic workflow (Figure 1) on pooled *D. magna* samples, we were able to identify 3532 unique phosphorylation sites in 1329 phosphoproteins.

**Figure 1.**
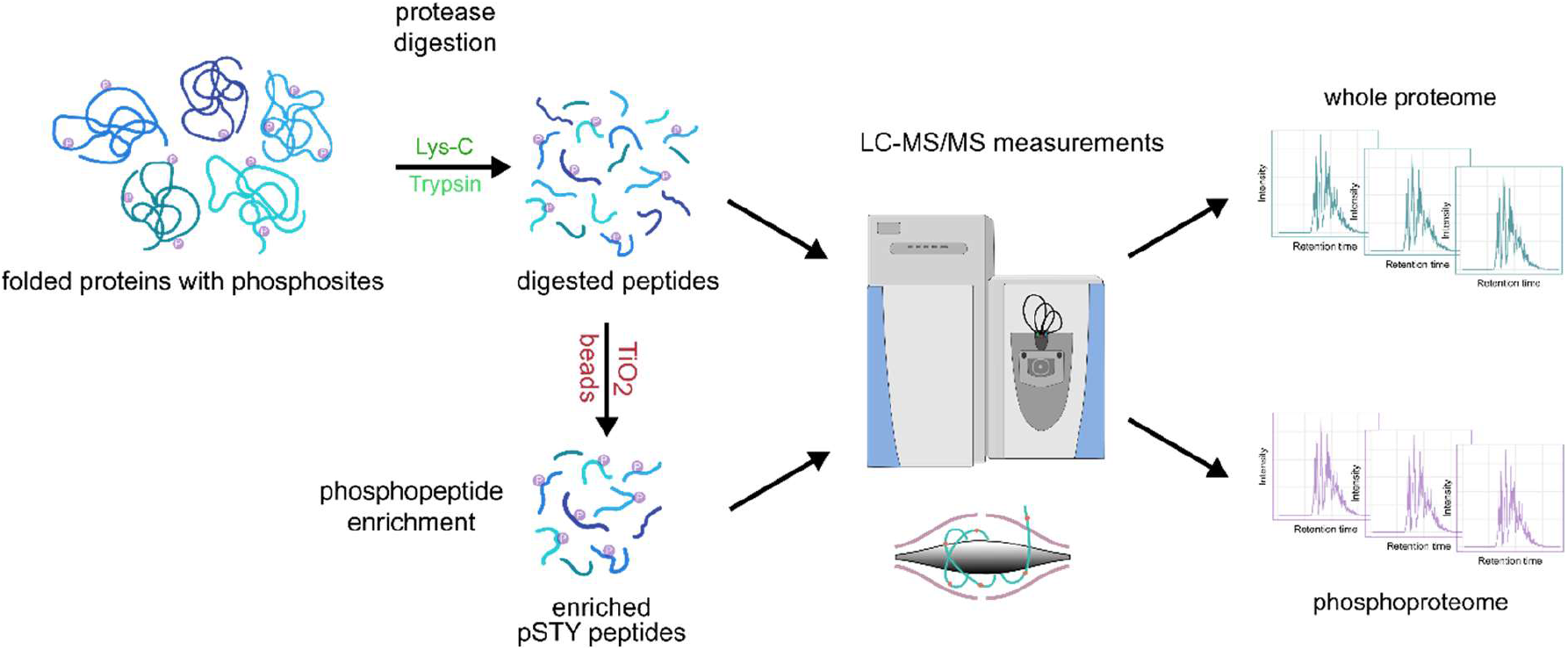
Phosphoproteomic workflow: After lysis, proteins were digested to subsequently enrich their phosphopeptides (pSTY refers to phosphorylation on serine, threonine, and tyrosine) before being measured on a mass spectrometer.

For the first time, we demonstrate a phosphoproteomic LC-MS/MS approach in combination with an in-solution digestion for *Daphnia magna*, providing an easy-to-perform yet effective method. To obtain sufficient material for phosphoproteomic analysis, 10 adult females of the *D. magna* clone K_3_4J were pooled in a reaction tube, the remaining water was carefully removed with filter paper and then snap-frozen in liquid nitrogen. The pooled samples (n=5) were homogenized by ultrasonication in 200 µl of lysis buffer (8 M urea, 50 mM ammonium bicarbonate) supplemented with protease inhibitors (one Ultra tablet mini per 10 ml buffer (Roche, Germany)). The protein quantification was done using the Pierce 660 nm Protein Assay (Thermo Fisher Scientific, U.S.A). After reduction and alkylation, sequential digestion, first with Lys-C (4 h, 37 °C) followed by trypsin (overnight, 37 °C), was performed. To obtain optimal conditions for phosphopeptide enrichment, a desalting step was done using the Sep-Pak® Kit (Sep-Pak Vac 1cc (100 mg) tC18 Cartridges, Waters, U.S.A) following the manufacturer’s instructions. For the enrichment of phosphopeptides the High-Select™ TiO_2_ Phosphopeptide Enrichment Kit (Thermo Fischer Scientific, U.S.A.) was used. In a final step, enriched phosphopeptides were dried using a vacuum concentrator (Bachofer, Germany).

LC-MS/MS analysis was performed using an Ultimate 3000 RSLC (Thermo Fisher Scientific, U.S.A.) connected to a Q Exactive HF-X mass spectrometer (Thermo Fisher Scientific, U.S.A.). Phospho-enriched peptides were resuspended in 15 µl 0.1% formic acid (FA), loaded on a trap column (PEP-Map100 C18, 75 µm × 2cm, 3 µm particles (Thermo Fisher Scientific, USA)) and separated on a reversed-phase column (PepMap RSLC C18, 75µm × 50 cm, 2µm particles, Thermo Scientific, U.S.A) at a flow rate of 250 nl/min. A 30-min gradient of 3-25% solvent B followed by 5-min increase to 40% was used. Solvent A consisted of 0.1 % FA, and solvent B of 0.1 % FA in ACN. After separation, the column was washed using 85% solvent B for 10-min. For MS/MS analysis, the data-dependent acquisition method consisted of cycles of one MS scan with a mass range of m/z 300-1600 at a resolution of 60000, followed by a maximum of 15 MS/MS scans at a resolution of 15000. We deposited the mass spectrometry proteomics data to the ProteomeXchange Consortium (http://proteomecentral.proteomexchange.org) via the PRIDE partner repository [19] with the project accession: PXD077473.

Raw mass spectrometric data was analyzed using MaxQuant [20] (v. 2.0.3.0) and Perseus [21] (v. 1.6.7.0). In MaxQuant, the Andromeda search engine score filter was left at the default setting of 40, and the enzyme specificity was set to trypsin. Cysteine carbamidomethylation was set as a fixed modification. N-acetylation of protein, methionine oxidation, and the phosphorylation of Serine, Threonine, Tyrosine (STY) were set as variable modifications. For protease digestion, up to two missed cleavages were allowed. The MS/MS spectra were matched against a *D. magna* UniProt FASTA database, which was run through CD-Hit [22] with a 95% sequence identity cut-off to reduce redundancy. In Perseus, phosphosites were filtered for a localization probability greater than 75%.

In total, we identified 3532 phosphorylation sites (Supplemental Table 1) and 3192 phosphopeptides, which could be assigned to 1329 phosphoproteins, with an FDR of 1%. Besides the study by Kwon et al. [15], in which they reported the first global screening of phosphoproteins in *Daphnia pulex*, this is, to our knowledge, the second phosphoproteome dataset in *Daphnia* and one of the most comprehensive phosphoproteomic analyses of aquatic organisms in recent years. Most of the phosphoproteomics studies on higher developed aquatic organisms in the last few years have focused on fish [23–26] while invertebrates [27, 28] have been studied to a lesser extent. Kwon et al. [15] identified 103 phosphorylation sites in 91 proteins in *D. pulex*. We compared the results of our study with the phosphoproteins identified by Kwon et al. [15] by blasting the identified *D. pulex* phosphoproteins against our *D. magna* results. We considered phosphoproteins that had at least 90% query coverage and 90% identity to be shared in both studies. This applied to 39 phosphoproteins, while 52 were not in our dataset, and 1290 are novel identifications (Figure 2A). The intensities of identified phosphoproteins show a dynamic range larger than five orders of magnitude. Gerritsen et al. [29] stated that MS-based phosphoproteomics must be able to handle a large dynamic range of phosphorylations in order to identify and quantify ultra-low level, dynamic phosphorylation events, as well as phosphorylations in high-abundance proteins at high stoichiometry. Furthermore, our data showed that 33.4% of the identified proteins were phosphorylated at a single residue (444 proteins), whereas 66.6% (885 proteins) were phosphorylated at two or more residues (Figure 2B). In more detail, we identified 30.8% of the phosphoproteins having 4 or more phosphorylation sites (410 proteins) and only 2.6% having more than 14 sites (35 proteins). In comparison to a study on mice by Huttlin et al. [30], where 80% of proteins contained multiple phosphorylation sites, with 50% being phosphorylated on four or more residues and 10% on more than 14 sites, our findings indicate fewer phosphorylations per protein. Sebé-Pedrós et al. [31] suggest that the phosphosignaling machinery is more sophisticated in vertebrates than in early diverging organisms or non-animal taxa, based on the greater number of proteins with multiple phosphorylation sites. This suggests that the comparatively lower number observed in our study may reflect an inherently lower phosphorylation level in daphnids. The observed distribution of phosphosites among the S/T/Y residues, with serine being by far the most frequently phosphorylated amino acid (91%), followed by threonine (8%), and tyrosine (1%) being the least phosphorylated, is consistent with several studies [32–35] (Figure 2C).

**Figure 2.**
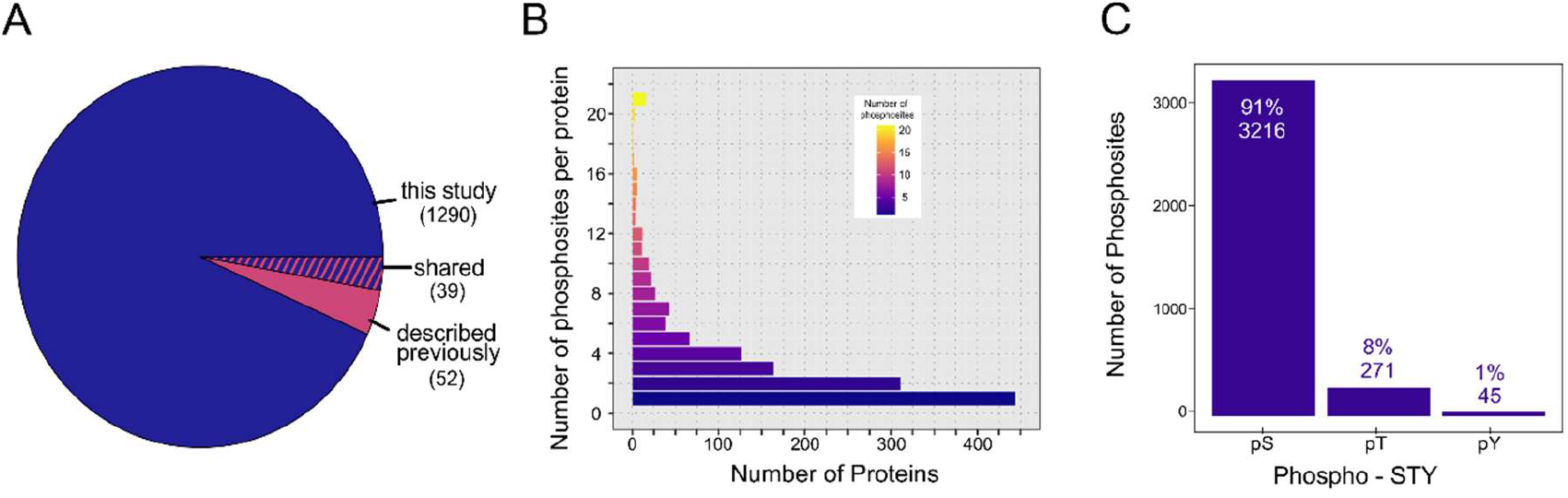
(A) The pie chart depicts phosphoproteins that were previously reported by Kwon et al. 2014 Kwon, Sim, Yun, Kim and Lee [15] and those that were identified in the present study, as well as those that were identified in both studies. (B) Distribution of the phosphorylation sites per protein over all phosphoproteins. Proteins with 21 or more phosphosites per protein were summarized. (C) Distribution of phosphosites across serine, threonine, and tyrosine residues.

To gain further insight into the biological functions of the identified phosphoproteins, a functional classification analysis was performed in PANTHER [36, 37]. For this, the corresponding *D. magna* sequences were blasted against *D. pulex*, using Blast+ [38] from NCBI, and were subsequently filtered for best hits. This output was used in PANTHER and plotted as a treemap using the “treemap” package in R Studio [39] (version 2022.07.2). The three most prominent gene ontology terms in the GO aspect “biological process” are cellular process (45.9%, GO:0009987), metabolic process (29.3%, GO:0008152), and biological regulation (19.9%, GO:0065007) (Figure 3A). For the GO aspect “molecular function”, two GO terms, namely binding (32.0%, GO:0005488) and catalytic activity (23.2%, GO:0003824), are the most represented (Figure 3B). The proteomic landscape is generally highly diverse in different organisms, as stated by Müller et al. [40]. Our findings are consistent with their discovery that proteins involved in metabolic processes constitute one of the most prevalent categories of proteins across all organisms. Furthermore, Humphrey et al. [41] pointed out that phosphorylation plays a crucial role in cellular metabolism, acting as a key switch that links metabolic enzymes into complex signal transduction networks and controls protein function through two main mechanisms: modulation of protein-protein interactions and modification of protein conformation.

**Figure 3.**
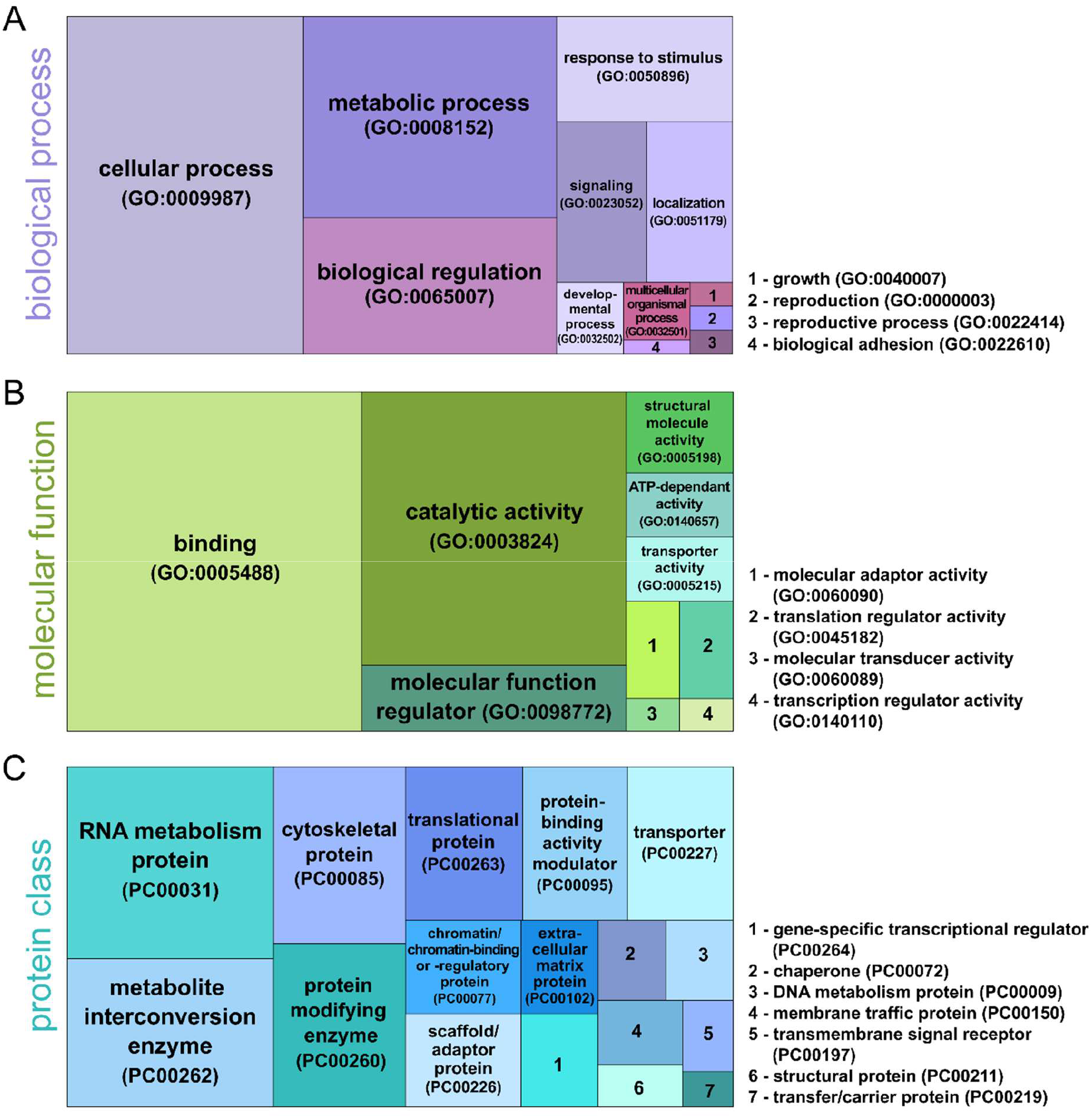
Treemap of gene ontology (GO) terms for phosphorylated proteins. (A) for the GO aspect “biological process” (B) for the GO aspect “molecular function” (C) for protein classes.

We also compared the results of our GO analysis with those of Kwon et al. [15] for *D. pulex*. Expectably, similar patterns are apparent in several sections. Binding and catalytic activity are also the most prominent GO terms in molecular function. Additionally, we found metabolic process, biological regulation, and response to stimulus to be one of the four main categories in the GO category biological processes. Our most abundant GO term, “cellular process”, is not present in Kwon et al. [15], which could possibly be the result of the higher phosphoproteomic depth of our dataset.

Finally, we used PANTHER to categorize the identified phosphoproteins according to protein classes. Here, we found phosphoproteins assigned to different protein classes, with fundamental RNA metabolism proteins (12.2%, PC00031) and metabolite interconversion enzymes (9.4%, PC00262) being the most abundant ones (Figure 3C). Also, the protein classes cytoskeletal protein (7.2%, PC00085), protein-modifying enzyme (6.6%, PC00260), and translational protein (5.5%, PC00263) were found.

In conclusion, our *D. magna* phosphoproteomic dataset delivers a comprehensive dataset of phosphorylation sites and their corresponding proteins, providing a robust foundation for elucidating regulatory phosphorylation mechanisms in this ecologically important freshwater species. This information is crucial for future studies aimed at deciphering the organism’s response mechanisms to contaminants as well as to environmental changes. The findings from this study are pivotal in advancing our comprehension of ecotoxicological dynamics and in reinforcing the central role of *D. magna* in ecological research and conservation efforts of freshwater ecosystems.

## ACKNOWLEDGMENTS

This study was funded by the Deutsche Forschungsgemeinschaft - Project number 391977956 - SFB 1357 to TF and C.L.

## DATA AVAILABILITY STATEMENT

The mass spectrometry proteomics data have been deposited to the PRIDE repository, dataset identifier PXD077473. Project Name: Phosphoproteomics in *Daphnia magna* as a tool to decipher molecular mechanisms in ecotoxicological studies.

